# Engineering, Generation and Preliminary Characterization of a Humanized Porcine Sickle Cell Disease Animal Model

**DOI:** 10.1101/2020.09.15.291864

**Authors:** Tobias M. Franks, Sharie J. Haugabook, Elizabeth A. Ottinger, Meghan S. Vermillion, Kevin M. Pawlik, Tim M. Townes, Christopher S. Rogers

**Affiliations:** Precigen Exemplar, 2656 Crosspark Rd. STE 100, Coralville, IA 52241, USA; National Center for Advancing Translational Sciences, Division of Preclinical Innovation, Therapeutics Development Branch, National Institutes of Health, Bethesda, MD 20892, USA; Lovelace Biomedical, 2425 Ridgecrest Dr. SE, Albuquerque, NM 87108, USA; UAB Schools of Medicine and Dentistry, Department of Biochemistry and Molecular Genetics, Birmingham, Al 35294, USA

**Keywords:** Sickle cell disease, hemoglobin, fetal beta globin, alpha globin, beta globin, gamma globin, Yucatan, pig, swine, porcine, animal model, CRISPR, Cas9

## Abstract

Mouse models of sickle cell disease (SCD) that faithfully switch from fetal to adult hemoglobin (Hb) have been important research tools that accelerated advancement towards treatments and cures for SCD. Red blood cells (RBCs) in these animals sickled *in vivo*, occluded small vessels in many organs and resulted in severe anemia like in human patients. SCD mouse models have been valuable in advancing clinical translation of some therapeutics and providing a better understanding of the pathophysiology of SCD. However, mouse models vary greatly from humans in their anatomy and physiology and therefore have limited application for certain translational efforts to transition from the bench to bedside. These differences create the need for a higher order animal model to continue the advancement of efforts in not only understanding relevant underlying pathophysiology, but also the translational aspects necessary for the development of better therapeutics to treat or cure SCD. Here we describe the development of a humanized porcine sickle cell model that like the SCD mice, expresses human ɑ-, β− and γ-globin genes under the control of the respective endogenous porcine locus control regions (LCR). We also describe our initial characterization of the SCD pigs and plans to make this model available to the broader research community.

## Introduction

Hemoglobin (Hb) is a protein tetramer consisting of two α-globin and two β-globin subunits that, when associated with a heme molecule, functions to efficiently recruit oxygen to tissues via RBCs [1-3]. In humans, multiple variants of β-like globin genes including embryonic (*HBE*; ε-globin), fetal (*HBG*; γ-globin) and adult (*HBB*; β-globin) genes are temporally regulated throughout development under the control of a super-enhancer designated a LCR [4-6]. The LCR elegantly recruits transcription modifying enzymes to promoters upstream of γ-globin and β-globin transcription start sites to modulate gene expression in accordance with proper developmental timing. For example, during fetal development, γ-globin is highly expressed and β-globin expression remains low. γ-globin protein complexes with α-globin and the fetal hemoglobin tetramer (HbF; ɑ_2_γ_2_) is produced. At birth, HbF represents approximately 90% of total hemoglobin. After birth, γ-globin expression decreases, and β-globin expression steadily increases until HbA (ɑ_2_β_2_) represents 98% of total hemoglobin at approximately one year of life. In a subset of people of African, Hispanic, Arabic, Mediterranean and Asian origin, an A→T mutation in the sixth codon of the β-globin gene codes for an amino acid switch from a polar glutamic acid residue to a hydrophobic valine residue (E6V) resulting in a structurally defective hemoglobin designated HbS (ɑ_2_β^S^_2_). Initially, newborns are clinically healthy due to the continued expression of γ-globin and presence of HbF; however, these children begin to show the symptoms of SCD after γ-globin protein levels decrease to below 20% of total hemoglobin and β^S^-globin protein production increases to levels above 80% within 9 months of birth. The HbS tetramers polymerize to form 14 stranded fibers that cause RBCs to become rigid and inflexible, occluding small capillaries in multiple organs. Sickle RBCs also become fragile and lyse which results in severe anemia and inflammation [7].

To better understand SCD, two groups have made knockout mouse models which faithfully recapitulate important hallmarks of the disease [8-10]. In one such “humanized” model, created by the Townes laboratory, the two mouse *Hba* genes were replaced with human *HBA* genes and the two mouse *Hbb* genes were replaced with a gene cassette containing human *HBG* and sickle human *HBB* (*HBB*^S^) along with a significant amount of upstream and downstream sequences that are important to retain temporal expression controlled by the LCR [8]. Humanized sickle cell mice are born healthy but experience a dramatic HbF to HbS transition soon after birth and subsequently begin to develop characteristics of SCD at approximately 3 weeks old [8]. These SCD characteristics include RBC’s with distinct sickle cell morphology and subsequent changes in blood cell composition; low RBC counts and anemia, organ pathology and shortened life span. Since production of the knockout transgenic sickle mice in 1997 [8, 9] and the humanized knock-in sickle mice in 2006 [10], these SCD mouse models have led to many critical advances in our understanding of SCD including the development of potential therapeutics [10-13]. Indeed, several groups to date have developed successful methods to correct SCD in the Townes mouse with the introduction of non-sickling β-globin into adult mice [11] or genetic replacement of *HBB*^S^ with *HBB* in blood cell progenitors [10, 13]. However SCD mice do present some limitations in modeling sickle cell lung disease, vasculopathy, bone marrow infarcts, bone marrow emboli, and retinopathy [14]. Therefore, the need exists for a large animal model of the disease that is directly relevant to human pathology to evaluate potential therapeutics. One promising model candidate, the swine, is more closely related to humans than mice and has similar cardiovascular, pulmonary, digestive, nervous and immune systems [15]. Additionally, several genetically engineered swine models have been generated for other diseases, setting precedence for use as a model for human genetic disorders [16, 17]. Swine also present opportunities in terms of the use of imaging and other diagnostic technologies to capture unique manifestations of disease as well as the ability to collect significantly large volumes of blood longitudinally in the same animal [18].

Here we created a humanized SCD or WT swine model in the Yucatan mini pig that is genetically equivalent to the Townes knock-in sickle cell mouse. We substituted porcine *HBA* (pα) with human *HBA* (hα) and porcine *HBB* (pβ) with a gene cassette containing human *HBG*-human *HBB*^*S*^ (hγ-hβ^S^) or human *HBG*-human *HBB* (hγ-hβ) with upstream and downstream sequences to allow temporal regulation by the endogenous pig LCR. Using somatic cell nuclear transfer (SCNT), we were able to birth healthy animals. Humanized SCD (HbS) pigs or corresponding WT (HbA) counterparts express α-globin, γ-globin, β^s^-globin, or β-globin proteins and form the expected HbA, HbS and HbF tetramers *in vivo*. Notably, RBC staining of blood from humanized 3-month HbS animals shows signs of characteristic sickle cell morphology as compared to control blood stains. Moreover, RBCs from HbS animals sickle dramatically under reduced oxygen tension, but RBCs from HbA animals do not sickle. We are currently observing and characterizing these animals in a 24-month natural history study for outward manifestation of the SCD phenotype as they age. We intend to make HbS and HbA animals available for future SCD research.

## Results

### Replacement of the porcine *HBA* and *HBB* alleles with the human *HBA, HBB* and *HBG* alleles

As a first step to generate hαγβ^S^ and hαγβ cells, we replaced both alleles of porcine *pHBA* with human *hHBA*. Wild-type (WT) pig fetal fibroblasts (PFF) were transfected with a plasmid (pFB-hα-Blast) containing the *hHBA*-Blast sequence flanked on the 5’ and 3’ ends by ∼ 1.3kb *pHBA* 5’ end (Exon 1) of the *pHBA* gene. Following 10-14 days of selection, surviving cells in each well of the 96-well plates were analyzed by PCR screening to detect the presence of correctly targeted hα-Blast. We used PCR primers that span the 3’ homologous recombination junction that should only be found in targeted cells (**Figure 1A, blue arrows**). Multiple colonies that appeared to be homozygous for the hα-Blast insertion (**Figure 1B**) were subjected to Blast and hα Southern blot analysis (**Figure 1C and 1D**). A correctly targeted cell line with no random integration was chosen for SCNT. Following embryonic transfer (ET) and successful implantation into an ovulating surrogate gilt, a fetal harvest was conducted at development day 35-40. Fetal cells were characterized to confirm correct targeting and frozen in preparation for future targeting to replace *pHBB* with *hHBB*^*s*^ or *hHBB*.

**Figure 1.**
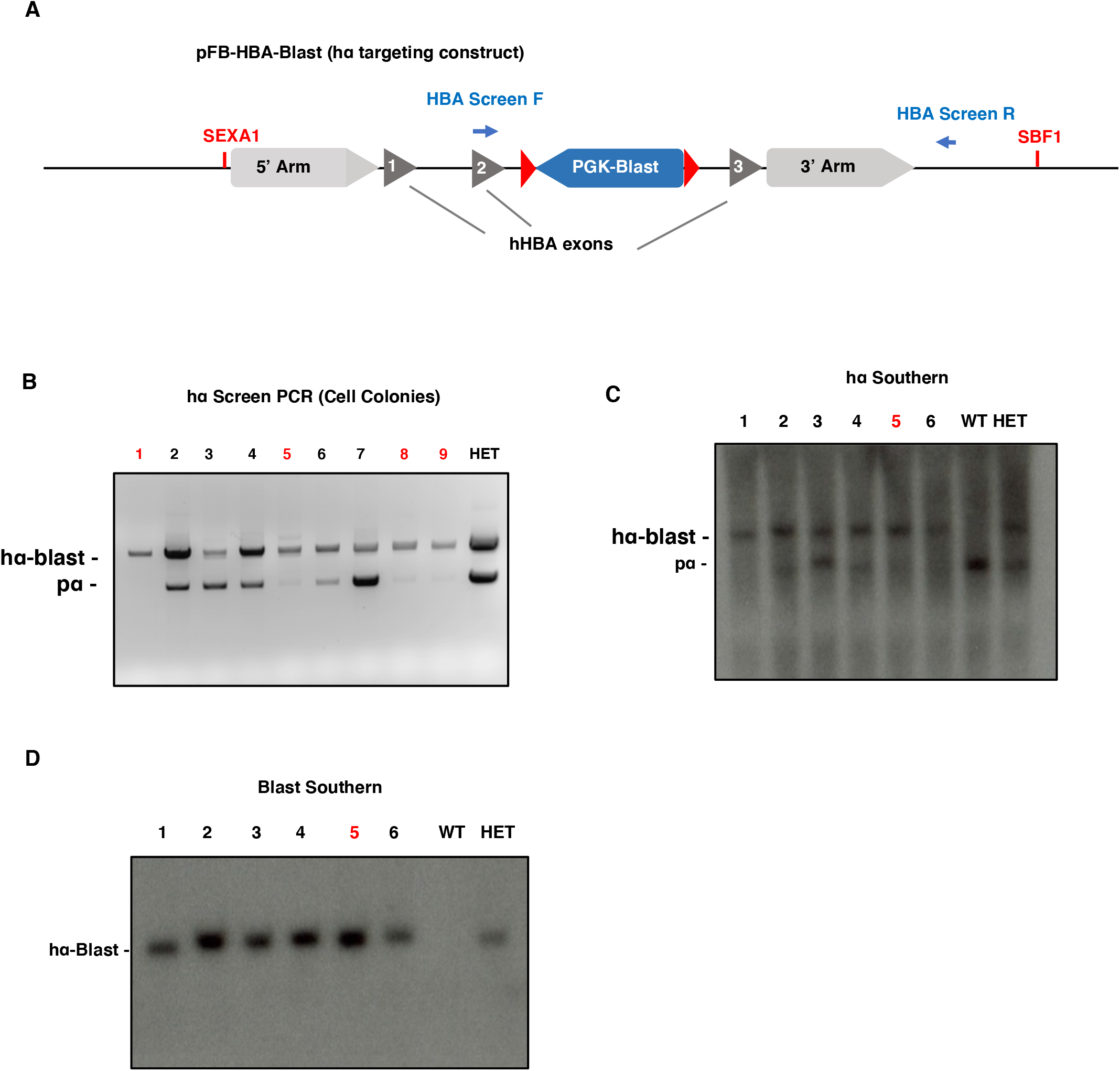
hα cell line targeting and characterization. (A) The *hHBA* gene was cloned into the pFB vector with a PGK-Blast resistance cassette (blue) in intron 2. *pHBA* 5’ and 3’ homology arms were cloned upstream and downstream of the *hHBA* gene, respectively (light grey). PCR primers to detect correctly integrated *hHBA* recognize the second exon of both *pHBA* and *hHBA* (“HBA Screen F”) and *pHBA* sequence downstream of the 3’ homology arm (“HBA Screen R”). (B) A representative PCR screen result showing identification of homozygote *hHBA* insertion into the *pHBA* locus (red font). (C) A representative *HBA* Southern blot showing identification of promising cell lines that have both *pHBA* alleles replaced with *hHBA*. (**Figure 1A**). The sample in lane 5 was used for SCNT and cloning to generate cells for hγ-Neo-β^S^ targeting. (D) A representative Blast Southern blot showing identification of promising cell lines that do not have random integration.

Next, we designed two targeting constructs, hγ-Neo-hβ^S^ or hγ-Neo-hβ (not shown) gene cassettes to replace both *pHBB* alleles with *hHBB*^*s*^ or *hHBB*. The *hHBG* portion of the hγ-Neo-hβ^S^ or hγ-Neo-hβ targeting vectors is comprised of three exons of *hHBG* and two introns as well as 1312bp and 2556bp of upstream and downstream sequence, respectively (**Figure 2A**). *hHGB* was fused with 2500bp of porcine sequence located on the 5’ end of *pHBB* exon 1 which served as the 5’ homology arm and the Neo resistance cassette which allowed for selection of correctly targeted cells. The *hHBB* portion of the targeting construct comprises the three exons and two introns of the *hHBB* gene as well as 818bp and 1819bp of upstream and downstream non-coding sequence, respectively (**Figure 2A**). The *hHBB* sequence was fused with 2500bp of porcine sequence located on the 3’ end of *pHBB* exon 3, which will serve as the 3’ homology arm (**Figure 2A**). To generate the sickle β-globin gene, a single base pair mutation corresponding to the human pathogenic SCD variant was made which changes the coding of amino acid 6 from glutamic acid to valine (**Figure 2A**). A guide RNA (gRNA) was generated that targets a region of *pHBB* that is not present in either the hγ-Neo-hβ^S^ or hγ-Neo-hβ targeting vectors. This ensures that once *pHBB* is replaced with *hHBB* or its sickle variant, using either targeting vector, Cas9 cannot re-target the allele.

**Figure 2.**
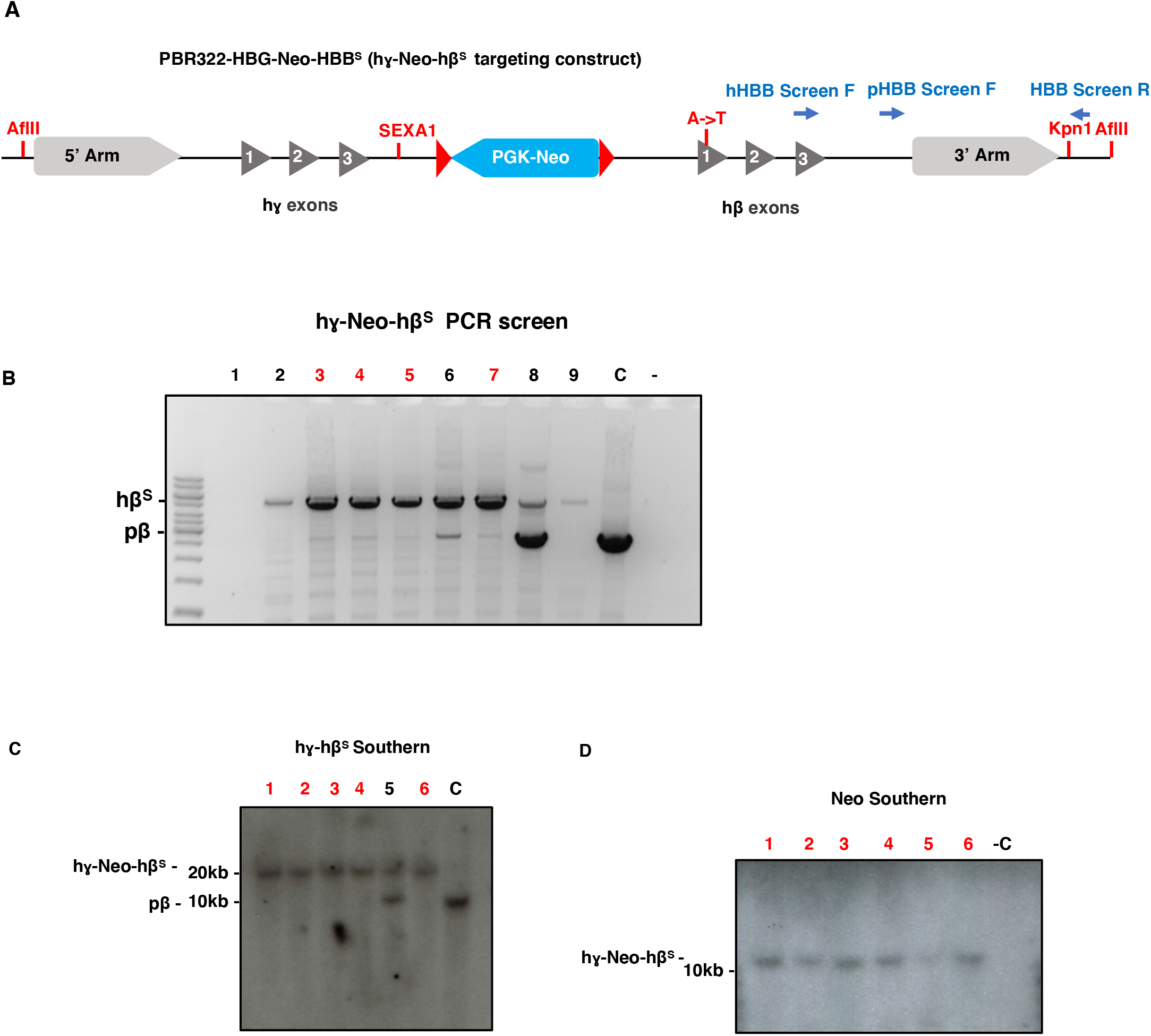
hγ-Neo-β^S^ cell line targeting and characterization. (A) The *hHBG* (along with upstream and downstream sequence), Neo cassette and *hHBB* genes (along with upstream and downstream sequence) were cloned into the pBR322 vector. An A→T substitution was made in the 6^th^ codon of *hHBB* which codes for the E6V SCD mutation (red). PCR primers to detect correctly integrated hγ-Neo-β^S^ recognize the third exon of *hHBB* (“hHBB Screen F”) downstream of the 3’ homology arm (“HBB Screen R”). Primers “pHBB Screen F” and “HBB Screen R” detect the presence of the *pHBB* allele. (B) A representative PCR screen result showing identification of homozygote hγ-Neo-β^S^ insertion into the *pHBB* locus. (C) A representative *HBB* Southern blot showing identification of promising cell lines that have both *pHBB* alleles replaced with hγ-Neo-β^S^ where gDNA was digested with AFLII (**Figure 2A**). (D) A representative Neo Southern blot showing identification of promising cell lines that do not have random integration from which one of the lines in red font was used for subsequent SCNT and cloning.

*hHBA* PFF cells (hαpβ) were then transfected with the hγ-Neo-hβ^S^ targeting plasmid and the Cas9/gRNA expressing plasmid, split into 96 well plates, and selected with PFF media containing Neomycin. After 14 days of cell growth and selection, cell colonies were assayed for correct targeting with PCR using a reverse primer that anneals to sequence outside the *pHBB* 3’ homology arm and two forward primers that recognize *pHBB* or *hHBB* sequence (**Figure 2A**). After this analysis, we determined that multiple colonies appeared to have both copies of *pHBB* replaced with either hγ-Neo-hβ^S^ (**Figure 2B**) or hγ-Neo**-**hβ (**data not shown**). We then used Southern blotting to test colonies that appeared to be homozygous for hγ-Neo-hβ^S^ for correct size and the presence of the Neo cassette (**Figure 2C and 2D**). Once several cell lines passed our quality control assays, they were treated with Cre recombinase to excise the Blast and Neo cassettes. These hαγβ^s^ or hαγβ PFF cells were then frozen in preparation for cloning and animal production.

### Generation of hαγβ^S^ and hαγβ knock-in pigs

To produce and characterize hαγβ^S^ and hαγβ pigs and assess their importance as a disease model, we chose to produce 15 animals of each genotype. Surrogate gilts were successfully implanted and gave birth to healthy piglets. All animals were genotyped using PCR with the same primers that were used to screen for respective hαγβ (not shown) or hαγβ^S^ colonies (**Figure 3A and 3B**). As shown in **Figure 3B**, all animals have the expected *hHBB*^*S*^ genotype. To confirm the efficient excision of Neo and Blast selection cassettes, we used a PCR assay with primers that flank the Neo cassette in hγ**-**hβ^S^ or the blast cassette in hα. This assay confirmed that both cassettes had been successfully excised (Figure 3C and 3D). To further assess the integrity of the *hHBG****-****hHBB*^*S*^ insertion, we conducted Southern blots. Genomic DNA (gDNA) samples from hγ**-**hβ^S^ piglets were digested with the restriction enzyme, AFLII which cuts outside the homology arms used to target *hHBG****-****hHBB*^*S*^ (**Figure 4A**). As seen in **Figure 4B**, all animals showed the correct *hHBG****-****hHBB*^*S*^ at ∼ 18kb while our non-humanized control *pHBB* band was only ∼ 8kb as expected. Similar results were observed with hγ**-**hβ animals (data not shown). To sequence the large ∼ 18kb region that was manipulated we conducted long range PCR (**Figure 4D**) to amplify 3 overlapping fragments. DNA bands were purified and sequenced to confirm the integrity of the hγ**-**hβ^S^ cassette and the presence or absence of the A→T mutation in hγ**-**hβ^S^ and hγ**-**hβ animals, respectively (**Figure 4C**). We conclude that we have successfully produced hγ**-**hβ^S^ and hγ**-**hβ animals that are suitable for characterization.

**Figure 3.**
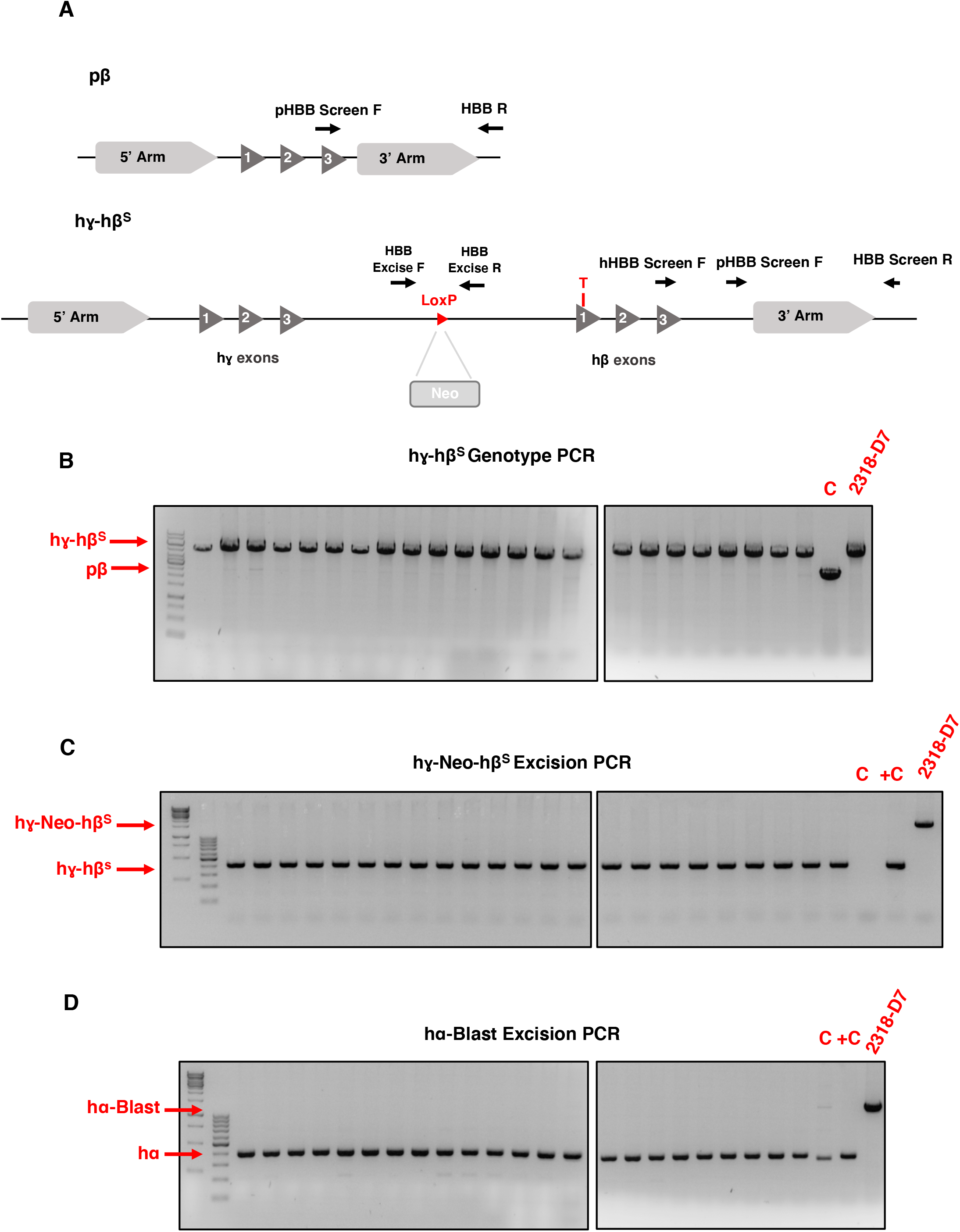
Genotyping and cassette excision of hγ-Neo-β^S^ targeted newborn animals. (A) Schematic of primers used to assess hγ-Neo-β^S^ genotype and Neo cassette excision. (B) hγ-Neo-β^S^ genotype PCR using 3 PCR primers (*pHBB* Exon 3F, hHBB Exon 2F and HBB R) to assess presence of hγ-β^S^ gene cassette and absence of *pHBB*. C lane = untargeted negative control, 2318-D7 = DNA from the 2318-D7 (cell line used to clone animals being genotyped). (C) Neo cassette excision was assessed using primers that flank the position of the cassette. (D) Blast cassette excision was assessed using primers (See primers section “HBA Geno F and HBA Geno R”) that flank the position of the cassette in the 2^nd^ intron of *HBA*.

**Figure 4.**
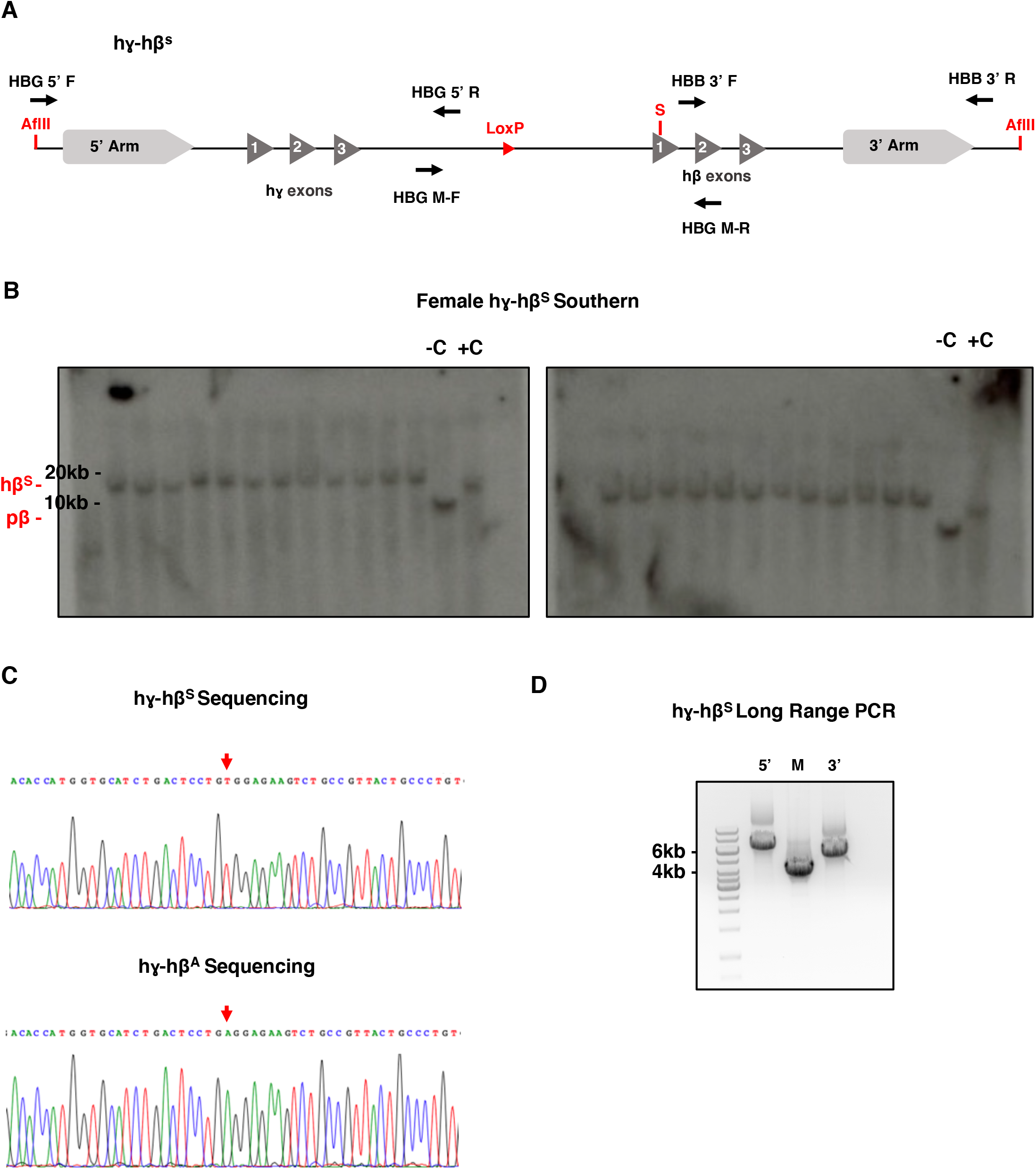
Southern blot and sequencing of hγ-hβ^S^ cassette in female hγ-hβ^S^ piglets. (A) Schematic of hγ-β^S^ gene cassette insertion. AFLII cut sites are outside the 5’ and 3’ *pHBB* homology arms. Primers used to amplify 3 overlapping fragments for sequencing are shown. (B) Southern Blot showing the replacement of *pHBB* (8kb) with the hγ-β^S^ (18kb) gene cassette. An untargeted cell line (-C) and previously characterized hγ-β^S^ (+C) cell lines are shown. (C) DNA sequencing result showing the presence or absence of A→T mutation that results in E6V amino acid substitution in hγ-β^S^ and hγ-βcell lines, respectively. (D) Long range PCR fragments purified to sequence the hγ-hβ^S^ gene cassette. 5’, Middle and 3’ fragments of the cassette are shown.

### Confirmation of *hHBA, hHBG* and *hHBB*^*S*^ genes and respective protein expression in knock-in pigs

Next, we wanted to confirm the expression of *hHBA, hHBG* and *hHBB*^*S*^ genes and respective protein products using reverse transcriptase PCR (RT-PCR) and western blotting techniques, respectively. As shown in **Figure 5A**, RT-PCR results indicate that *pHBA* and *pHBB* gene expression is absent in the humanized hαγβ^s^ model, but present in our non-humanized control indicating that pig globin subunit expression has been replaced with human globin subunit expression in the porcine hαγβ^S^ SCD model (hαγβ not shown). To further confirm that α-globin, γ-globin and β-globin proteins are being expressed in the hαγβ^S^ model, we extracted protein from the blood of five hαγβ^S^ animals at birth and 6 months of age and conducted western blot analysis with antibodies that detect human α-globin, γ-globin and β-globin proteins, but not pig α-globin and β-globin proteins (pigs do not endogenously express the *HBG* gene). As shown in **Figure 5B**, all hαγβ^S^ animals express α-globin, γ-globin, and β-globin proteins, but the control non-humanized animal does not. Importantly, we still observe high levels of γ-globin protein expression at 6 months post birth. If our model recapitulates the human disease, then it is expected that γ-globin expression should decrease at some point during the first year of life and be replaced by β-globin expression. We continue to monitor these animals at approximately 3-month intervals in a planned 24-month natural history study. The animals are currently 9 months old at the time of submission of this manuscript.

**Figure 5.**
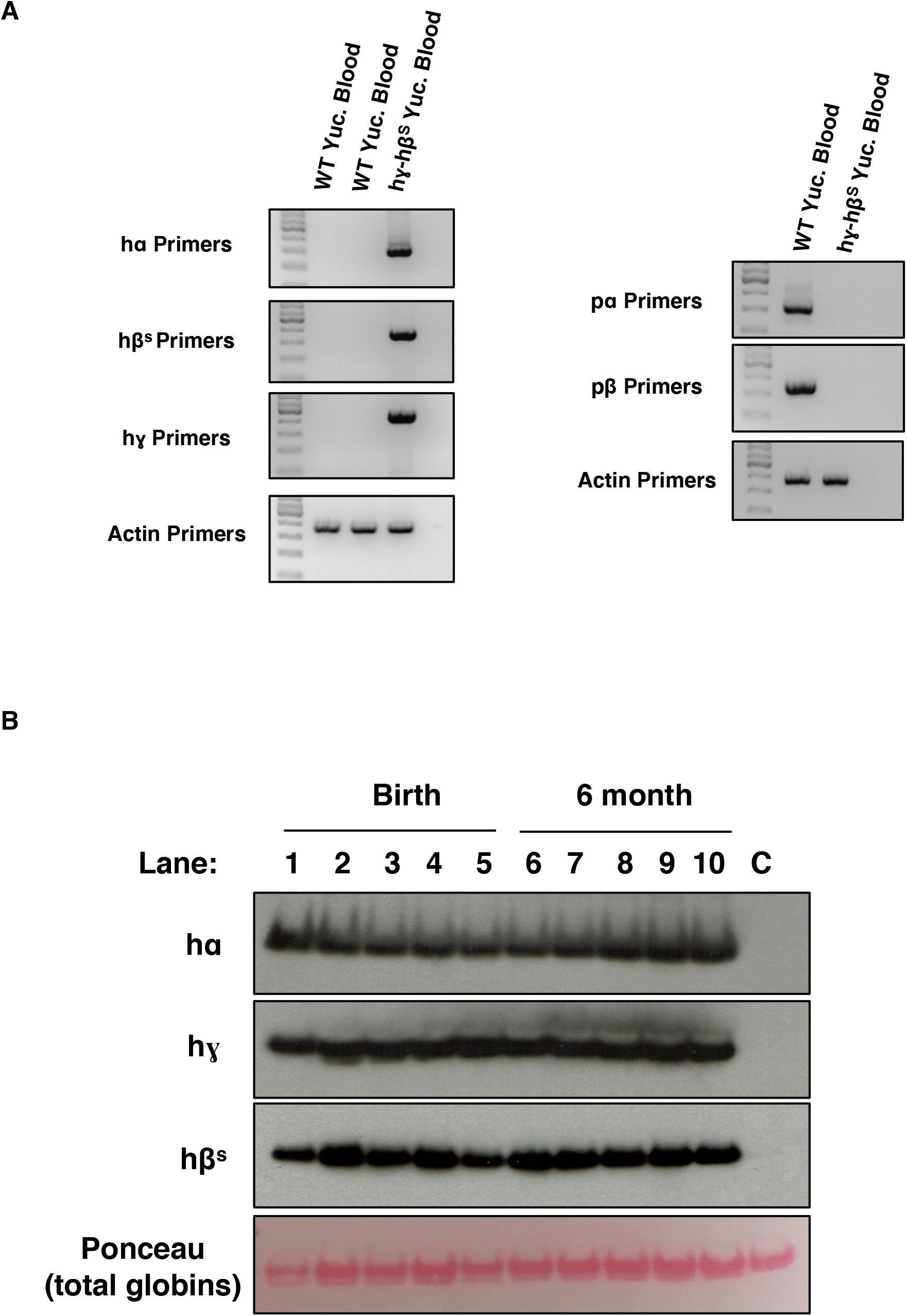
Human *HBA, HBG* and *HBB*^S^ RT-PCR and western blot of protein products. (A) RT-PCR showing expression of *hHBA, hHBB*^S^ and *hHBG* only in the blood of an hγ-β^S^ animal but not in the two WT animals (left figure) and *pHBA* and *pHBB* only in a WT animal but not a hαγβ^S^ animal (right figure). (B) Western blot showing protein expression of hα, hγ, and hβ^S^ in the blood of 6-month old hγ-β^S^ animals but not in a control (“C”) WT Yucatan animal. Total globin staining via ponceau is shown as a loading control.

### Hemoglobin complexes from humanized porcine SCD animals mimic human SCD patients

We next wanted to determine whether Hb from our porcine SCD animals forms similar complexes to those observed in human SCD patients. We conducted isoelectric focusing gel electrophoresis which allows us to separate different hemoglobin species (HbF, HbA, HbS, HbF acetylated) based on their differing migration through poly-acrylamide gels. If pig HbA is successfully substituted with human HbF and HbS or HbA then we expect to observe a predictable shift in Hb profile. Indeed, as observed in **Figure 6A**, hemoglobin extracted from hαγβ (“AA”) and hαγβ^S^ (“SS”) models partition similarly to hemoglobin extracted from cord blood of normal (“AA”) and SCD patients (“SS”), respectively. In contrast, WT non-humanized pig HbA runs higher and in a single band.

**Figure 6.**
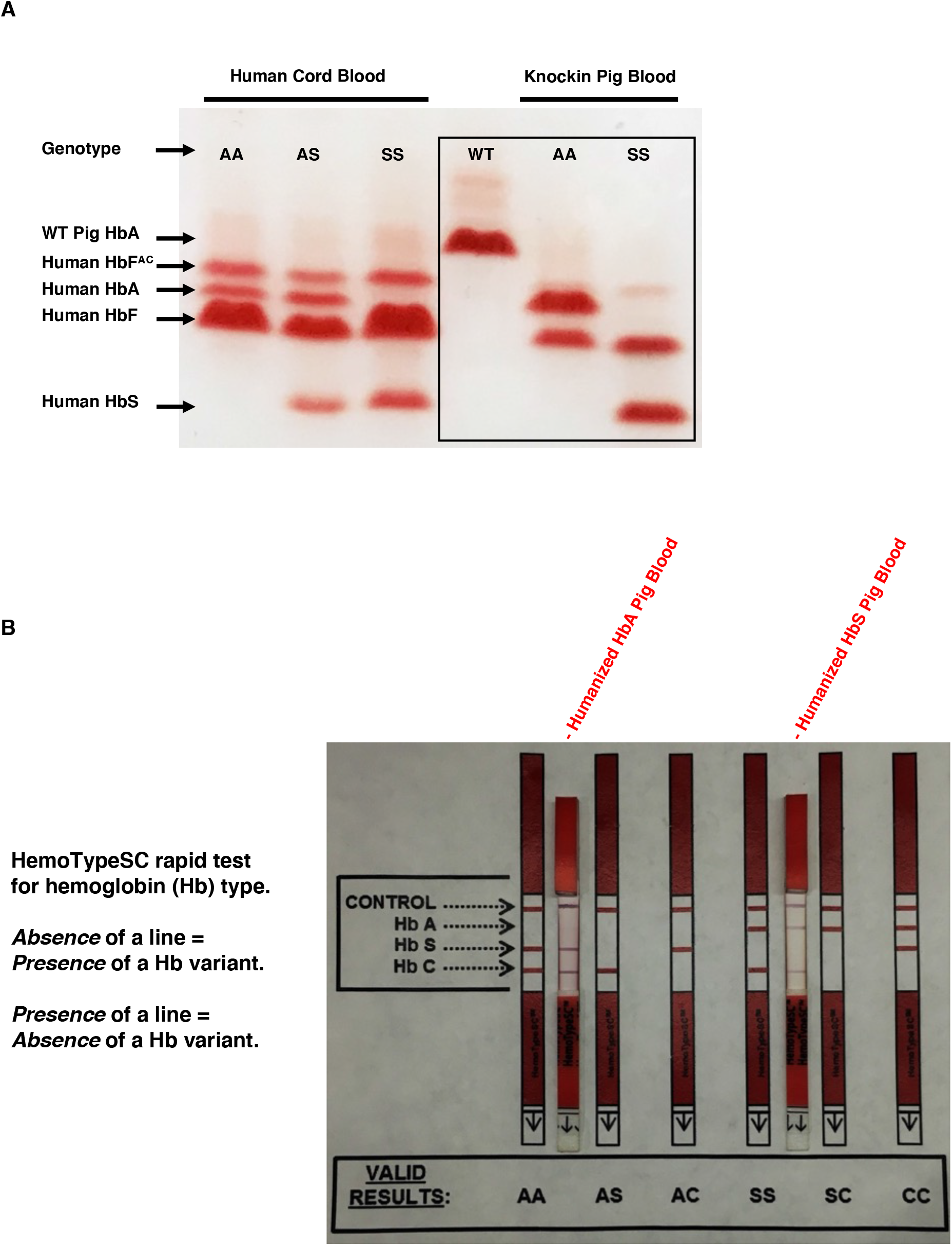
Hb from humanized pigs behaves similarly to human Hb. (A) Isoelectric focusing of hemoglobin species from human patients (left side) or non-humanized (WT) and humanized pigs (right side). For human cord blood or humanized pigs, AA = Normal Hb, AS = sickle cell trait Hb, SS = Sickle Hb. (B) HemoTypeSC™assay. The absence of a band indicates the presence of that Hb variant.

If Hb in our SCD model is forming complexes like those that trigger pathogenicity in humans, then we expected that a diagnostic test for SCD will correctly identify HbA and HbS pigs like in human patients. HemoTypeSC™is a clinical test commonly used to diagnose patients with SCD. This test uses monoclonal antibodies to block the appearance of a red band next to the indicated Hb species. Thus, the absence of a band on the test indicates the presence of the Hb variant in question. As shown in **Figure 6B**, the HemoTypeSC™test correctly diagnoses our HbA and HbS models. We conclude that functional human HbA, HbS and HbF are present in our porcine models and form complexes that behave similarly to HbA, HbS and HbF from human blood.

### HPLC and Mass Spectrometry confirm presence of HbF, HbA and HbS in humanized pigs

The isoelectric focusing experiments described above suggest that human Hb is forming the expected HbF, HbA and HbS complexes in our humanized pigs *in vivo*. To further confirm these results, we conducted standard high-performance liquid chromatography (HPLC) and reverse phase HPLC (RP-HPLC). As shown in **Figure 7A**, HbF, HbA and HbS complexes elute as expected in HbA and HbS samples. In contrast, WT pigs that do not express humanized Hb have only one HbA peak that elutes, but at a different retention time than human HbA. RP-HPLC experiments, which separate Hb into individual monomers, indicate the presence of human ɑ-, γ- and β^S^- or β-globin or only pig α- and β-globin in non-engineered WT Yucatan pigs as expected **(Figure 7B)**. Notably, two peaks at distinct retention times were observed for the α-globin subunit. One allele of *hHBA* (protein subunit designated hɑ^*^ in the chromatogram) varies from the expected hɑ by 1 amino acid at position 131. This occurred because the last 50 nucleotides of the *pHBA* allele match the *hHBA* allele except for 4 nucleotides at positions 393-396 in the coding region of *pHBA* DNA sequence. During targeting of the *hHBA* allele in the female PFF cell lines, crossing over occurred before the 4 non-matching nucleotides on one allele instead of after as was expected to occur. This 4-nucleotide substitution resulted in a single amino acid S131N substitution variant. We are currently generating HbS and HbA males using targeted male PFF lines devoid of the S131N variant. Subsequent breeding campaigns will generate litters with the correct amino acid at codon 131. As a result, future generations will express identical hα proteins and the 2^nd^ peak in the HPLC will not exist. Importantly, as it pertains to the current studies, all the analyses to date suggest that Hb tetramers are forming normally even in the presence of hα^*^. Moreover, we predict that the hα^*^ protein will have little if any effect on the phenotype of HbA and HbS pigs.

**Figure 7.**
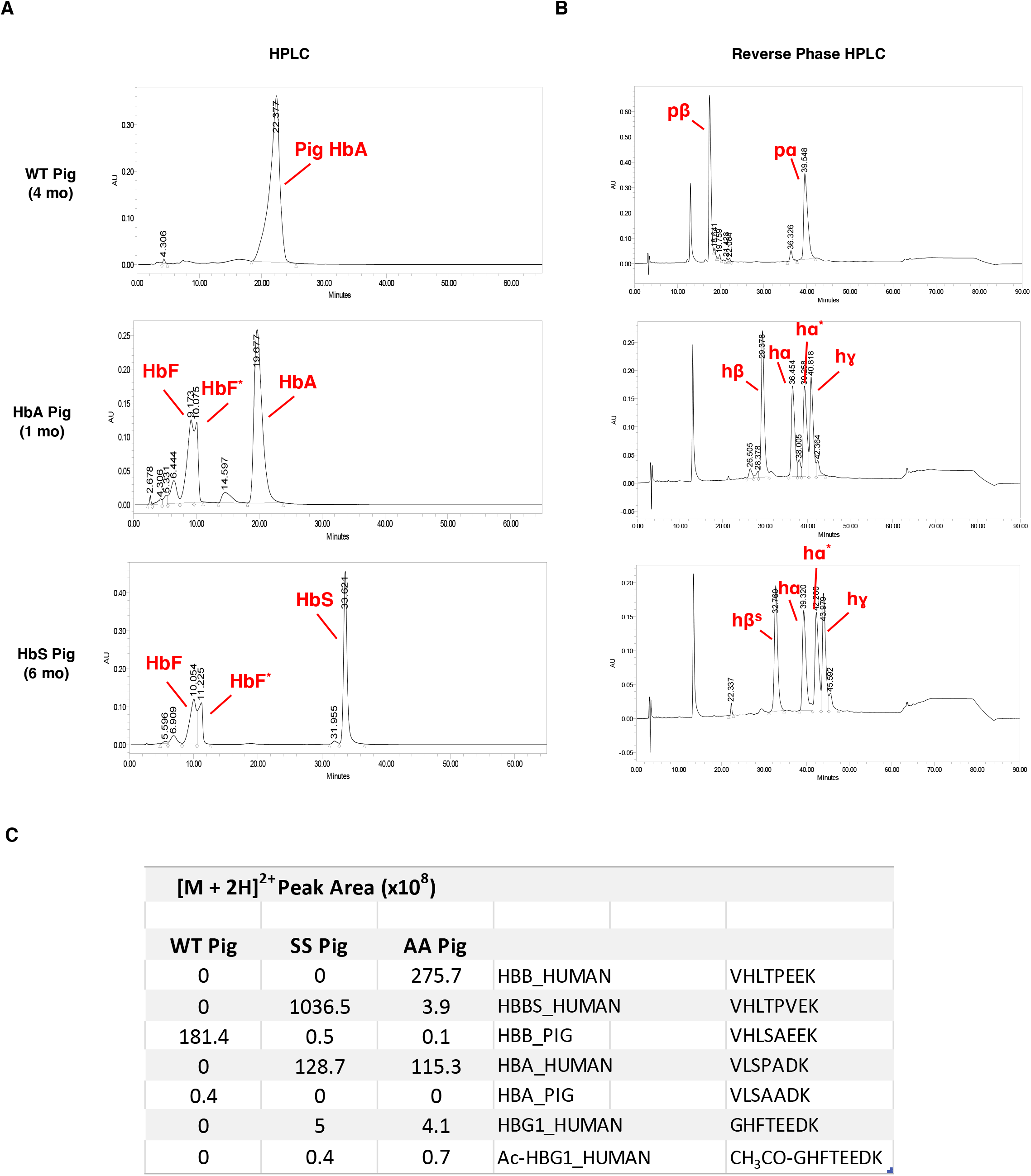
HPLC with blood from WT, HbA and HbS pigs. (A) HPLC assays showing the presence of only HbA in a control non-humanized WT pig and HbF and HbA or HbS in humanized pigs. HbF and HbF^*^ differ in that HbF tetramers contain hα, and HbF^*^ tetramers contain hα^*^ (see below) (B) Reverse phase HPLC showing the presence of individual globin monomers, hα, hγ and hβ^S^ or hβ. hα and hα^*^ differ in that they have a serine and an asparagine at amino acid position 131, respectively. (C) Quantification of globin peptides identified by mass spec analysis. Some contaminating peptides are observed in HbA (HBB^S^ human and HBB pig) and HbS (HBB pig) samples. This is a result from cross contamination of blood or protein during the mass spec sample preparation process which is typical for this type of analysis.

Subsequent mass spectrometric (MS) analysis in WT non-humanized Yucatan, HbA and HbS animals further indicate that HbA and HbS animals express only human globin subunits. Moreover, HbA pigs express hβ-globin while HbS animals express hβ^S^-globin **(Figure 7C)**. Although HbA and HbS pigs continue to highly express hγ-globin beyond 6 months of age we predict that we will observe the expected reduction in hγ-globin expression as animals approach 1 year of age. We conclude that the correct globin subunit proteins are expressed in our humanized pigs and that HbF, HbA and HbS complexes are forming correctly *in vivo*.

### Porcine SCD pigs have abnormal RBC morphology

Sickle shaped RBCs are the hallmark of SCD and are the root pathophysiological cause of SCD symptoms. To determine if our HbS pigs have sickle shaped RBCs, we prepared stained blood smears followed by light microscopy. At 3 months, sickle RBCs were occasionally observed in HbS animals but not in HbA animals or in wildtype non-humanized swine (**Figure 8A**). At 6 months of age, the sickled RBC phenotype was still relatively mild, presumably due to the persistence of HbF. We anticipate that the proportion of cells with sickle cell shapes will increase in HbS animals in the coming months. To further evaluate the sickling potential of the RBCs in the HbS pig, we next determined whether we could override the protective effect of HbF by treating RBCs with sodium metabisulfite (MBS) which decreases oxygen tension in RBCs and triggers HbS polymerization and cell sickling. As shown in **Figure 8B**, we observed significant sickling in MBS treated RBCs from HbS animals but no morphological changes in RBCs from WT pigs or humanized HbA pigs. These evaluations show that as early as 3 months of age, the HbS animals already demonstrate early characteristics of SCD with the predictive potential for full disease manifestation demonstrated by the outcome of the MBS assay.

**Figure 8.**
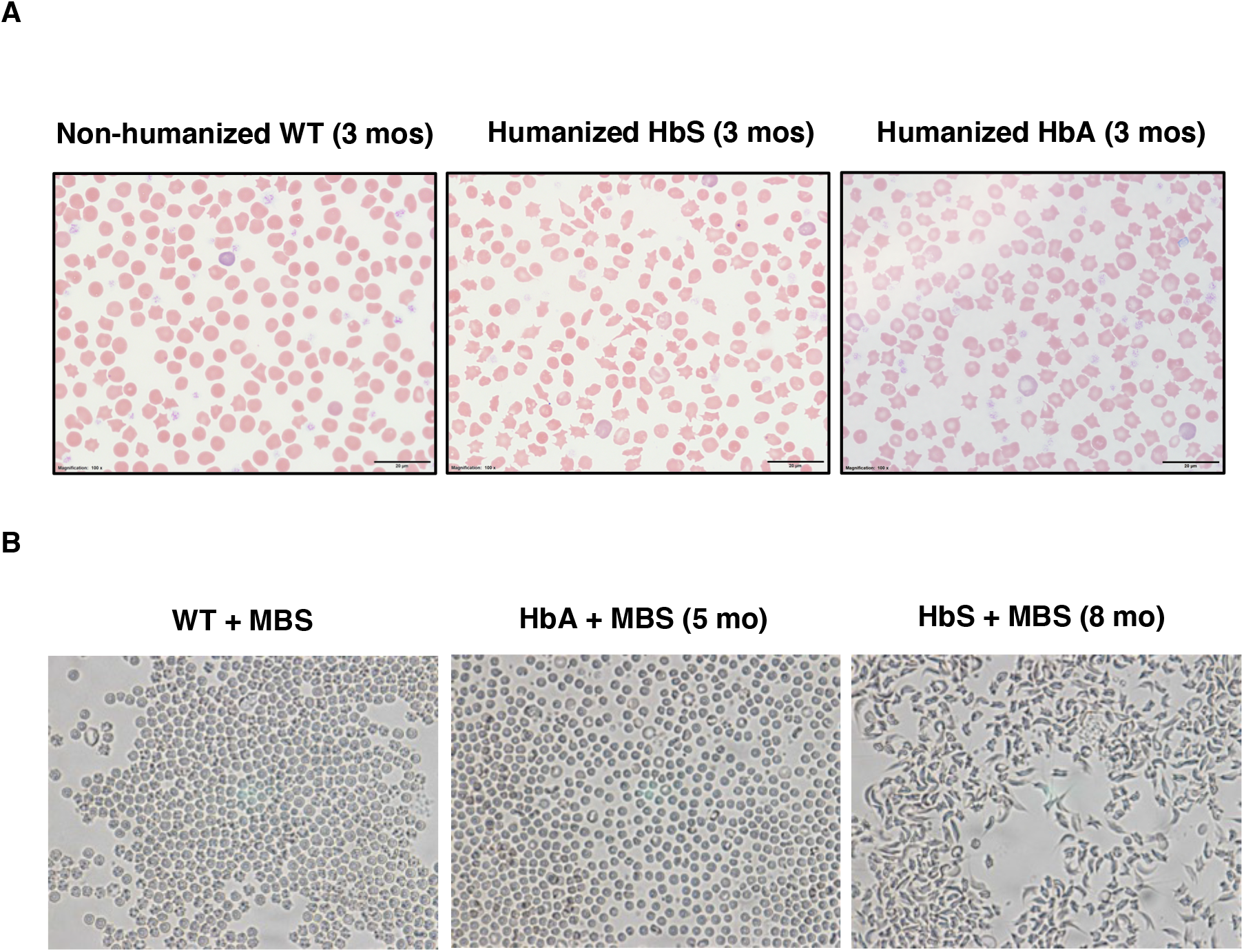
RBCs from HbS animals have sickle cell morphology. (A) Representative photomicrographs of peripheral blood smears showing blood cell morphology from WT (non-humanized), age-matched (i.e. 3 months) HbA and HbS pigs. (B) Light microscope images of RBCs after MBS treatment from WT, HbA (5-month old) and HbS (8-month old) pigs.

## Discussion

Approximately 300,000 infants are born worldwide each year with SCD, with a disproportionate number of these infants being born in the developing world [19]. At least 89,000 people in the United States currently live with SCD [20]. Although over 90 percent of children in the developed world can expect to reach adulthood, the median age of survival for those with SCD is 48 years of age, more than 20 years less than their demographic peers [21]. Of those with SCD that live into adulthood, they face a chronic debilitating condition that often leads to disability and death [22]. Furthermore, annual healthcare costs for SCD patients in the United States exceed $1 billion dollars annually [22].

Therapeutics have generally been very limited for patients with SCD. Hydroxyurea has been proven to be somewhat beneficial for pediatric patients and has helped to improve pediatric outcomes [23]. The only currently approved available clinical cure for SCD comes from allogeneic bone marrow transplantation (alloBMT) and has been performed in approximately 2000 patients in the developed world [22]. The regimen for this treatment is intense, finding donors is difficult and adults are often precluded. Currently, promising myeloablative gene therapy and genome editing clinical trials are ongoing [22]. However, since most individuals with SCD live in the developing world, the limitations for diagnosis and treatment due to infrastructure, logistics, and economics is significant and widespread use of gene therapy will not be available. Thus, a small molecule drug still has the most potential for world-wide benefit.

This SCD pig offers a unique animal model platform where researchers will have the opportunity to study in greater detail, the temporal and longitudinal manifestation of this devastating illness in tissues, organs, and systems like the human patient. Additionally, a pig model of SCD may also provide the potential to better predict the bench to bedside translational aspects of a given therapeutic under development that a mouse model may not be fully capable of realizing. One obvious tissue that is highlighted for this disease and a large animal model is the ability to collect and examine blood samples from the same pigs multiple times over an extended period to understand the potential long-term benefits of these therapeutics in a model more similar to humans than mice. Another prominent aspect of SCD research is pain associated with the sickling of RBCs resulting in excruciatingly painful vaso-occlusive crisis events. Gigliuto et. al., concludes that “pigs represent the most suitable model for evaluating pain because of their similarities to human in immunoreactivity in peptide nerve fibers.” In addition, “changes in distribution and axonal excitability of nociceptive and non-nociceptive fiber classes” are also stated as similar in human and pigs [24].

Less than five years ago, Hydroxyurea was the only approved drug for SCD and since then, while there have been several new approvals to target different aspects of SCD symptomology, to date, no additional disease-modifying therapeutic has been approved that directly targets the underlying biology, e.g., capitalizing on the human genetic validated approach of activating fetal hemoglobin production. There have been and there are several such drugs under development, but none have made it to approval for safe human use. Perhaps the potential of a novel, more translatable large animal model as an additional preclinical research tool will help to increase the likelihood of better small molecule candidates getting into the clinic and achieving a higher probability of success in clinical safety evaluation and efficacy studies.

Our SCD animals are currently 9 months of age and are expected to begin to show outward signs of SCD by twelve months if proper switching of HbF to HbS occurs. We have confirmed robust expression of human ɑ-, γ- and β^S^ at one month and 6 months of age. Although γ-globin levels have yet to be fully replaced by β^s^- or β-globin, we have observed sickle cell RBC morphology, including sickle shaped RBCs in HbS animals. Currently, as part of a two-year natural history study (NHS), we are conducting a full characterization of the model, including HPLC analyses at ∼ 3 month intervals to monitor the expression of globin subunits, urinalysis, hematology, blood chemistry, electrophysiological, pain and related behavioral measurements, imaging of brain and lung, other cardiopulmonary diagnostic evaluations as well as gross and histopathological analyses for animals that die prematurely or make it to the end of the study period. If this model does recapitulate the human disease, we envision that it will be valuable for the broader research community as a translational research model to not only contribute more to the understanding of the pathophysiological manifestations of SCD, but also biomarker discovery, and as a translational bridge for bench-to-bedside studies supporting therapeutic development efforts between the existing mouse models that have been mainstays of SCD research and clinical trials in humans. We view the SCD pig as a transitional model between preclinical and clinical development of experimental therapeutics. We are committed to making available the data and results of the ongoing NHS study via additional publications and other public resources.

## Methods

### Engineering of gene targeting constructs for generation of humanized PFF cell lines for cloning of humanized pigs

WT pig fetal fibroblasts (PFF) were transfected (Lipofectamine LTX (Invitrogen)) with a plasmid containing the hα-Blast sequence (ensemble reference ENSG00000206172.8) flanked on the 5’ and 3’ ends by ∼ 1.3kb *pHBA* homology arms **(Figure 1A)** and a plasmid that expresses Cas9 and a guide RNA (gRNA) targeting the 5’ end (Exon 1) of the *pHBA* gene (See **Figure S1** for plasmid maps).

Next, we designed a targeting construct to replace both *pHBB* alleles with a hγ-Neo-β^S^ or hγ-Neo-β gene cassette (**Figure S1**). The *hHBG* portion of the targeting vector is comprised of three exons of *hHBG* and two introns as well as 1312bp and 2556bp of upstream and downstream sequence, respectively (**Figure 2A**). *hHBG* was fused with 2500bp of porcine sequence located on the 5’ end of *pHBB* exon 1 which served as the 5’ homology arm and the Neomycin resistance cassette which allowed for selection of correctly targeted cells. The *hHBB* portion of the targeting construct comprises the three exons and two introns of the *hHBB* gene as well as 818bp and 1819bp of upstream and downstream non-coding sequence, respectively (**Figure 2A**). The *hHBB* sequence was fused with 2500bp of porcine sequence, which will serve as the 3’ homology arm (**Figure 2A**). A single base pair mutation corresponding to the human pathogenic SCD variant was made using mutagenesis PCR. This variant changed the coding of amino acid 6 of *HBB* that is not present in the hγ-Neo-β^S^ or hγ-Neo-β (not shown) targeting vectors. This ensures that once *pHBB* is replaced with hγ-Neo-hβs Cas9 cannot re-target the allele.

### Gene targeting

Approximately 5×10^5^ WT pig fetal fibroblasts were transfected (Lipofectamine) with a plasmid containing either the hα-Blast **(Figure 1A)** or hγ-Neo-β (not shown) or hγ-Neo-β^S^ targeting constructs **(Figure 2A)** and a plasmid that expresses Cas9 and a guide RNA targeting the 5’ end of the *pHBA* or *pHBB* genes. Twenty-four hours after transfection, cells were transferred to a series of 96-well plates, and G418 was added to the media. Following 10-14 days of selection, surviving cells in each well of the 96-well plates were expanded by splitting among (approximately 3000 cells split to each well depending on confluency) three separate plates: 96-well culture plates for further cell expansion, 96-well culture plates for potential cryopreservation, and 96-well PCR plates for cell lysis to prepare lysates for PCR genotype screening to confirm the presence of correct gene targeting.

### Genotyping PCR

Primers (See below) indicated in blue (**Figure 1A and 2A**) were used to amplify the 3’ homologous recombination region of the correctly inserted targeting construct or a smaller fragment of the untargeted pig WT allele.

### Southern blot

For analysis of potential targeted colonies, gDNA was purified from cell colonies and 8 ug of DNA was digested with restriction enzymes overnight at 37°C (hα or Blast blots -Sbf1 and SEXA1, hγ-hβ^s^ Blot-AFLII, Neo Blot-Sbf1 and SEXA1. DNA samples were run on 0.8% agarose gel at 110v for 1.5 hrs. Next the gels were depurinated in 0.25% HCL, denatured in alkaline denaturing buffer and washed twice in alkaline transfer buffer. DNA was transferred to nitrocellulose membrane overnight at room temperature. Membranes were crosslinked using a stratalinker and incubated for 30 min in prewarmed blotting buffer. Blocked membranes were incubated for 2 hours with DNA probe synthesized using a Prime-a-Gene® kit (Promega) and washed in mild and stringent Southern wash buffers. Blots were exposed to X-ray film at −80°C for 6-48 hrs.

### Western blot

100 µL of whole blood was pelleted by rapid centrifugation and pellets were lysed in 100 µL of RIPA lysis buffer with proteinase inhibitor cocktail (Roche). Following an additional centrifugation step, 75 µL of supernatant was added to a new tube containing 75 µL of SDS sample buffer (Invitrogen). Concentrated samples containing SDS LB were diluted 1:10 in SDS LB to prevent over saturation of the western blot signal due to the extremely high globin concentrations in RBCs. 5 µL of 1M DTT was added and samples were denatured for 10 min at 70° C. 5 µL (for hα, hγ) or 0.5 µL (for hβ^s^ blot) of diluted lysate was loaded onto a gradient gel (4-12%) (Invitrogen). Gels were run at 130 volts for 1 h and transferred to PVDF membranes. Next membranes were exposed to primary antibodies for 30 min (Santa Cruz; alpha: sc-514378, beta: sc-21757, gamma: sc-21756). After 3 x 5-min washes, membranes were exposed to secondary antibodies for 30 min and washed an additional 3 times. Following treatment with chemiluminescent reagents to activate HRP-conjugated antibodies, blots were exposed to X-ray film for 1-60 seconds.

### Blood smear

Approximately 5 µL of whole anticoagulated blood from non-humanized WT, hγβ or hγβ^S^ animals was spotted onto a microscope slide and smeared using an angled spreader slide to create an evenly spread monolayer of RBCs. Slides were stained using a Siemens HemaTek automatic slide stainer with a modified Wright-Giemsa stain. Slides were dried and viewed by light microscopy, and representative photomicrographs were obtained from the ‘feathered edge’ of the smear.

### Hemoglobin typing using HemoTypeSC™

1 µL of whole blood/EDTA was added to 200 µL of water in a 5 mL Fluorescence-activated cell sorting (FACS) tube and mixed by pipetting. A HemoTypeSC™ Test Strip (Silver Lake Research Corp., Azusa, CA, www.hemotype.com) was immersed into the resulting hemolysate and left undisturbed during hemolysate migration up the strip. After 10 min, the test strip was removed and visually compared to a hemoglobin (Hb) variant reference sheet to detect human HbA and HbS.

### Hemoglobin typing by isoelectric focusing (IEF)

Hemoglobin typing was performed using a RESOLVE Hemoglobin Kit (PerkinElmer, Inc., Waltham, MA, www.perkinelmer.com) and a Multiphor II Electrophoresis System (GE Healthcare Life Sciences, Marlborough, MA, www.cytiva.com). Samples for Hb variant analysis were prepared by mixing 10 µL of whole blood/EDTA with 100 µL of RESOLVE Hb Elution Solution. IEF gel preparation and electrophoresis of 5 µL sample volumes was performed according to the RESOLVE Hemoglobin Kit instructions. After Hb variants completed migration to their isoelectric points, the positions of bands from pig blood samples and human cord blood samples were compared. WT pig HbA (adult, α_2_β_2_) and human HbF^AC^ (fetal, α_2_^AC^γ_2_), HbA (adult, α_2_β_2_), HbF (fetal, α_2_γ_2_) and HbS (sickle, α_2_β^s^_2_) were identified.

### Normal and reverse-phase HPLC

100µL of anticoagulated whole blood was added to 1250µL of normal saline in an Eppendorf tube. Samples were inverted several times to mix and spun at 14000 rpm for 2 minutes. Supernatant was removed and the cell pellet was washed in 1250µL of saline and spun again at 14k rpm. After supernatant removal, 1 mL of HPLC grade water and cells were vortexed to create a homogenous hemolysate. 500µL of this hemolysate was added to 500µL of HPLC grade water. 1 drop of 1% KCN solution was added. HPLC columns (Synchropak CM-300) were preconditioned prior to use with two column washes of HPLC grade H_2_O_DI_ through the column for 120 minutes at the flow rate of 1ml/minute. After H_2_O_DI_ wash, columns were equilibrated by running 70% working buffer A (Bis Tris 146.5g, ammonium acetate 4.62g, potassium cyanide 2g in 2L H_2_O_DI_) and 30% working buffer B (Bis Tris 146.5g, ammonium acetate 26g, potassium cyanide 2g, 408.2g sodium acetate in 2L H_2_O_DI_) for 60 minutes at 1ml/minute flow rate. Hemolysates are loaded onto the column and run. Different hemoglobin types are calculated based on the area under the curve as percentage. Minor peaks are calculated first and the major peak last making it a total of 100%.

For reverse phase HPLC a Supelco C5 wide-pore HPLC column was used on a Waters Alliance 32 HPLC system. Prior to loading hemolysates, each column was prepared and conditioned with developer A (20% acetonitrile, 80% H_2_O_DI_, 0.07% TFA) and developer B (60% acetonitrile, 80% H2O_DI_, 0.07% TFA).

### LC-MS/MS of globin polypeptides

Hemolysates were diluted in 50 mM Tris-HCl, pH 8.0, and the protein concentrations were determined using Qubit fluorometry (Invitrogen). 20 µg of each sample was processed by SDS-PAGE using a 4-12% Bis-Tris NuPage mini-gel with the MOPS buffer system. The target region was excised from each lane and processed by in-gel digestion with trypsin using a ProGest robot (DigiLab). Briefly, bands were washed with 25 mM ammonium bicarbonate followed by acetonitrile, reduced with 10 mM dithiothreitol at 60°C followed by alkylation with 50 mM iodoacetamide at room temperature, digested with trypsin (Promega) at 37°C for 4h, and finally quenched with formic acid and the supernatant was analyzed directly without further processing. Half of each digested sample was analyzed by nano LC-MS/MS with a Waters NanoAcquity HPLC system interfaced to a ThermoFisher Q Exactive mass spectrometer. Peptides were loaded on a trapping column and eluted over a 75 µm analytical column at 350nL/min; both columns were packed with Luna C18 resin (Phenomenex). The mass spectrometer was operated in data-dependent mode, with the Orbitrap operating at 70,000 FWHM and 17,500 FWHM for MS and MS/MS, respectively. The fifteen most abundant ions were selected for MS/MS. Hb tryptic peptides were identified using Mascot (Matrix Science) with the following parameters; enzyme: Semi-Trypsin, Fixed modifications: Carbamidomethyl (C), Variable modifications: Acetyl (N-term), Deamidation (N,Q), Oxidation (M), Pyro-Glu (N-term Q), Mass values: Monoisotopic, Peptide Mass Tolerance: 10 ppm, Fragment Mass Tolerance: 0.02 Da, Max Missed Cleavages: 2.

### Red Blood Cell Sickling Test

7.5 µL of whole blood/EDTA were mixed on a microscope slide with 7.5 µL of a 2% sodium metabisulfite solution (0.2 g MBS/10 mL H_2_O). A cover slip was immediately applied, and the edges sealed with clear nail polish (Sally Hansen Hardener). After incubation at room temperature for 24 hours, slides were examined microscopically for cell morphology changes and sickling using 60X dry and 100X oil-immersion objectives.

### Study Approval

This study was carried out in accordance with the recommendations in the NIH *Guide for the Care and Use of Laboratory Animals*. All animals were developed and housed in the AAALAC-accredited facilities of Exemplar Genetics. Standard procedures for animal husbandry were used throughout. The IACUC of Exemplar Genetics approved all animal experiments.

## Oligonucleotides Used in Study

### HBA Primers

#### HBA Screen

HBA Screen F: GAC CCG GTC AAC TTC AAG GTG AG

HBA Screen R: CAG CAT CTC TAA TCT CGT TCA

#### HBA Genotype

HBA Geno F: CAA GAC CTA CTT CCC GCA CTT CG

HBA Geno R: CAG CAG GCA GTG GCT TAG GAG

#### HBA RT

hHBA RTF: CGC TGG CGA GTA TGG TGC G

hHBA RTR: CCG CAG GGG TGA ACT CGG

pHBA RTF: CAA GCT GGC GCA CAC GGC GC

pHBA RTR: GAA CGG AGG GGT TGA AAT CAT CGG

### HBG-HBB Primers

#### HBB Screen

hHBB Screen F: CAT CAG TGT GGA AGT CTC AGG ATC G

pHBB Screen F: CAG GAC AGA GCG CAC AGG AG

HBB Screen R: GCT TCA TCA TTC GCT AGT TCT CAT CTG G

#### HBB Genotype

HBB Excise F: GAA GTT GAC AAC TGC AAT GAT AAC CTG G

HBB Excise R: CAA TGT GCT CTG TGC ATT AGT TAC TTA TTA GG

### HBB RT Primers-

hHBB RT F: GGA GAA GTC TGC CGT TAC TGC

hHBB RT R: GGT GAA TTC TTT GCC AAA GTG ATG G

pHBB RT F: GAG AAG GAG GCC GTC CTC GG

pHBB RT R: GTC ATG GCC AAG GCG GCG A

#### hHBG-hHBB LR PCR Primers

HBG 5’ F: GGT ATC ATG CTT AGT GAA ATT AGT GAG ACA GG

HBG 5’ R: CAC AAA CAT TAG GTC CGT AAC ACC ATC

HBG Middle F: GAATCAGCAGAGGCTCACAAGTCAG

HBB Middle R: CCAAATAGTAATGTACTAGGCAGACTGTGTAAAG

HBB 3’ F: CATCAGTGTGGAAGTCTCAGGATCG

HBB 3’ R: GCTTCATCATTCGCTAGTTCTCATCTGG

#### hHBG RT Primers

hHBG RT F: GGT CAT TTC ACA GAG GAG GAC AAG G

hHBG RT R: GGT ATC TGG AGG ACA GGG CAC TG

#### Primers used to make gRNAs (Cloned into X335 vector-BBSI Digest)

pHBA gRNA F: CAC CGG GTG CTG TCT GCC GCC GAC A

pHBA gRNA R: AAA CTG TCG GCG GCA GAC AGC ACC C

pHBB gRNA F: CAC CGC ACC CTG TGG AAC CAC GCC C

pHBB gRNA R: AAA CGG GCG TGG TTC CAC AGG GTG C

#### hHBB direct seq

hHBB direct seq F: GCT GAG GGT TTG AAG TCC AAC TCC

hHBB direct seq R: GGA CTC AAA GAA CCT CTG GGT CC

## Acknowledgements

We would like to thank the following individuals for their contributions: Li-Chen (Jane) Wu and Chiao-Wang (Joe) Sun from the University of Alabama at Birmingham for the IEF gel in Fig. 6A. Michael Ford from MS Bioworks for LC-MS/MS data analysis and interpretation. Niren Patel and the Kutlar Laboratory at Augusta University for their assistance with the HPLC assays. Andrew Gigliotti and Vanessa Ortega Lovelace Biomedical Research Institute for their assistance with blood smear photomics and analyses. London Toney and Bryce Comstock from NCATS for their administrative assistance with the project management of this study. The Exemplar Genetics farm staff Frank Rohret, Judy Rohret, Jason Struzynski, Trisha Smit, Justin Van Kalsbeek and Cris Van Ginkel for their hard work and diligence handling the animals used in this study. Allison Ripperger at Exemplar Genetics for laboratory assistance.

## Support

This work was supported in part by the National institutes of Health (NIH) HEAL Initiative and in part by the Intramural Research Program of the National Center for Advancing Translational Sciences, NIH under Contract No. HHSN261200800001E from the National Cancer Institute, NIH. The content of this publication does not necessarily reflect the views or policies of the Department of Health and Human Services, nor does mention of trade names, commercial products, or organizations imply endorsement by the U.S. Government.

---

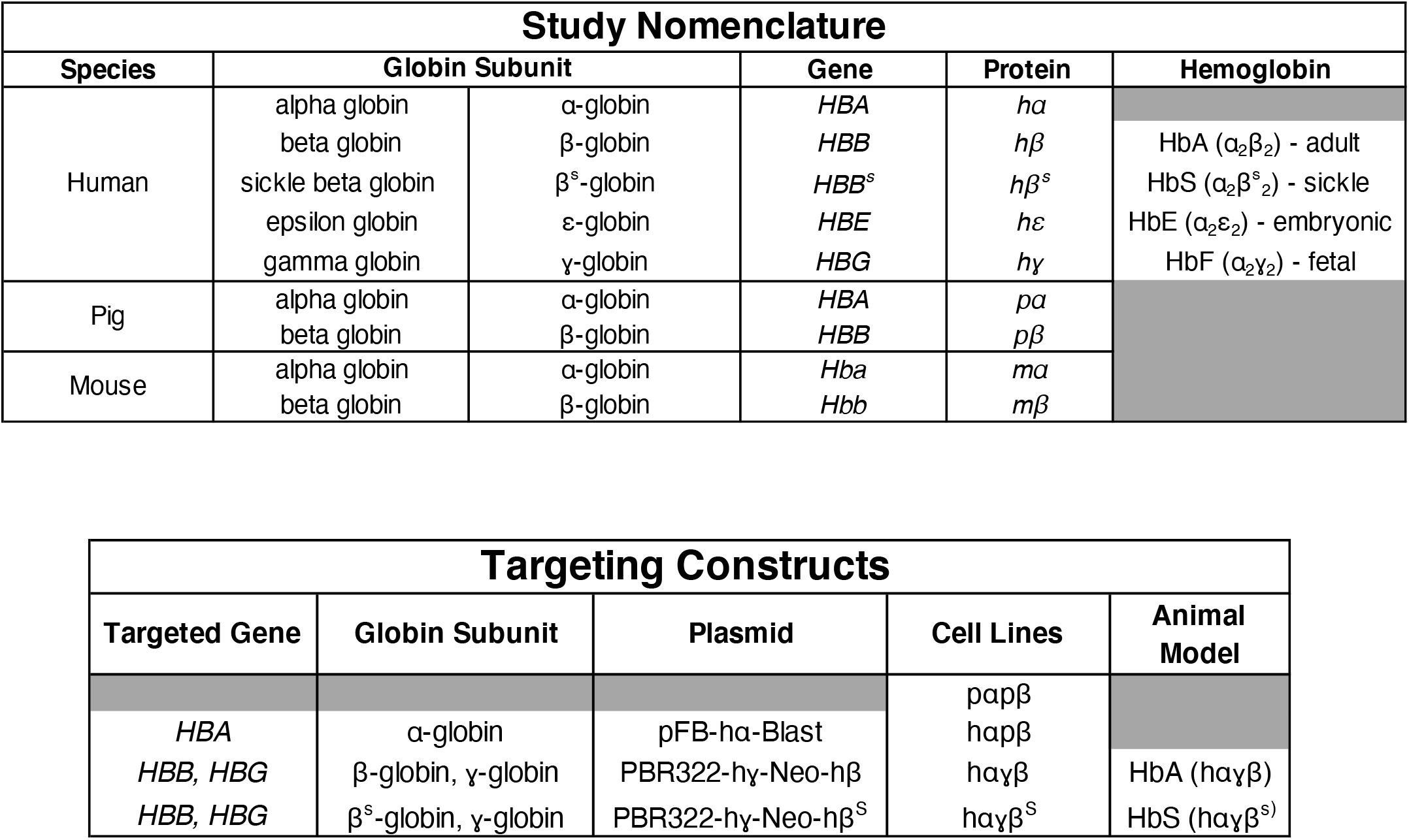

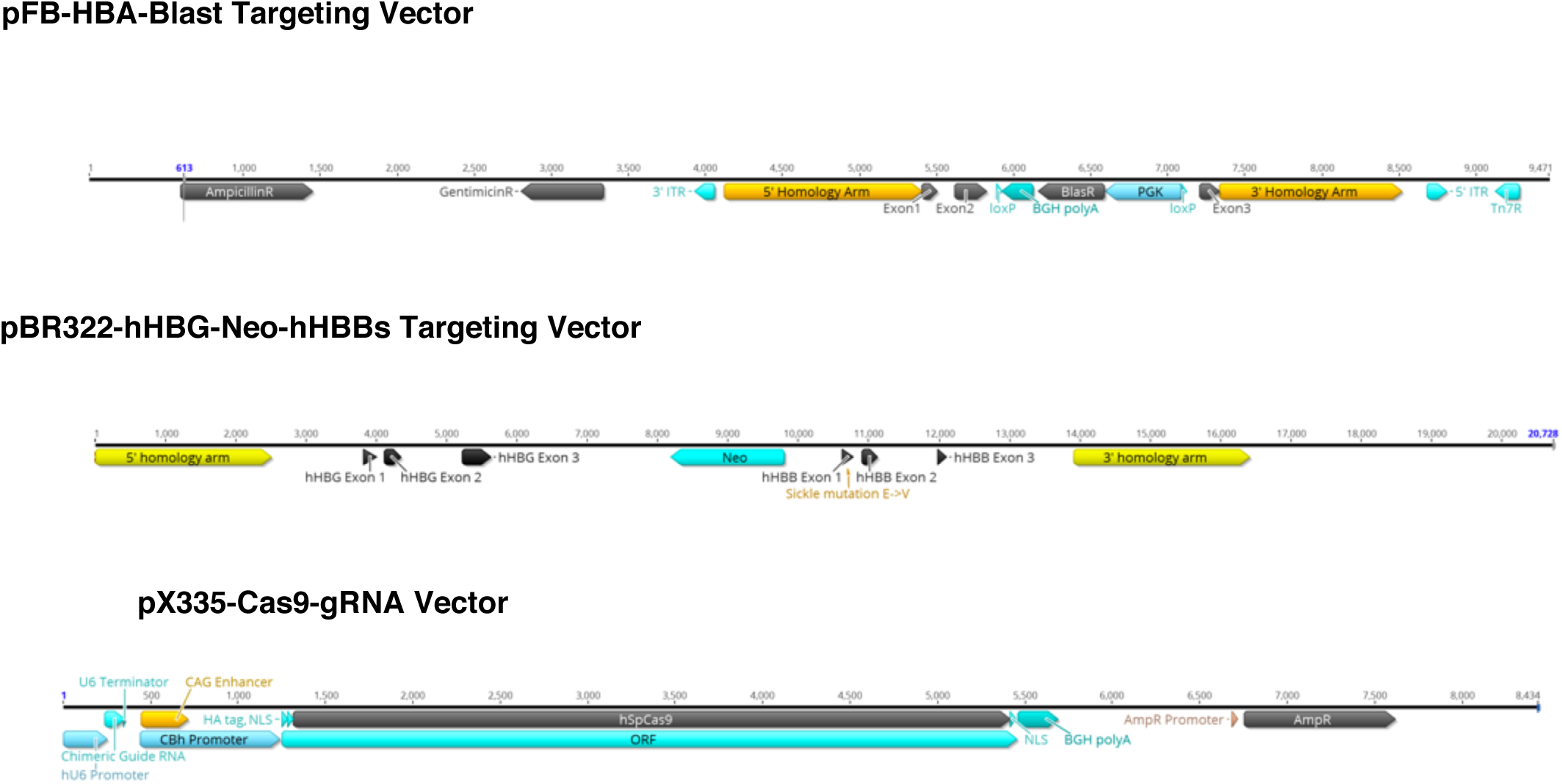

## Notes

### Competing Interest Statement

The authors have declared no competing interest.

